# Altered Social Cognition Associated with Kleptomanic and Instrumental Thefts

**DOI:** 10.64898/2026.08.19.745606

**Authors:** Yukiori Goto, Mehlika Iclal Cakir, Satoshi Yoshino, Chikara Kita, Moojun Won, Young-A Lee

**Author notes:** Correspondence: Yukiori Goto, Ph.D. Department of Artificial Intelligence and Technology Graduate School of Informatics Kyoto University Yoshida Honmachi, Sakyo-ku, Kyoto 606-8501, Japan.

## Abstract

Theft, including shoplifting, extorts a pervasive societal and economic burden. However, the neurobehavioral mechanisms underlying recurrent theft remain sparsely understood. In this study, we investigated social cognition deficits in theft recidivists with kleptomania (TR+K) and instrumental theft recidivists without kleptomania (TR-K) compared to control subjects without criminal records (CT), for which the Social Norms Questionnaire (SNQ-22) to assess explicit moral knowledge, alongside the Dictator Game (DG) and Hawk-Dove Game (HDG) to evaluate discretionary and competitive resource allocation with others, respectively, were administered. Bayesian statistical analyses revealed that all groups demonstrated comparable social norm recognition in SNQ-22 and prosociality in the DG. However, distinct behavioral profiles emerged in specific contexts, such that TR+K exhibited more unfairness than CT and TR-K at discretionary resource allocations in the DG, whereas in the HDG, TR-K demonstrated more aggressive, resource-monopolizing responses, particularly when against an aggressive opponent, than CT and TR+K. These results suggest that theft recidivism may stem from contextual failures rather than general deficits in moral knowledge, which are distinct between TR+K rooted in the internal factor, such as heightened loss aversion, and TR-K characterized by impulsivity over the external factor, such as social conflicts with others.

**SIGNIFICANCE STATEMENT:** Theft imposes a massive economic and societal burden, yet the psychological mechanisms driving repeat offenses remain poorly understood. We found that repeat offenders recognized social rules as well as non-offenders; therefore, recurrent theft was not caused by a lack of moral understanding. Instead, their criminal offenses are based on specific contextual failures in decision-making. In particular, kleptomania may be driven by internal factors such as loss aversion, while instrumental theft may be driven by impulsive reactions to external social conflicts. The findings suggest that shifting the focus from generalized punishment to context-specific psychological rehabilitation is important for reoffending prevention.

## INTRODUCTION

Theft, particularly in the form of shoplifting, represents a pervasive social problem that imposes a substantial financial burden on global economies (Blanco et al., 2008; Yaniv, 2009). Despite these severe economic and societal consequences, the neurobehavioral mechanisms underlying theft remain barely understood. This gap in the literature is particularly evident regarding recurrent theft and kleptomania, an impulse-control psychiatric disorder characterized by an inescapable urge to steal (Grant, 2006; Asaoka et al., 2025).

Forensic psychology and neurocriminology increasingly view criminal offenses, including theft, as fundamentally linked to breakdowns in sociality (Walters, 2022a, b). Moral disengagement, which is the psychological process by which people selectively deactivate their moral standards to justify self-serving behavior (Bandura, 1991), is thought to play a prominent role in theft, alongside contextual empathy deficits, which is an inability to generate empathetic concern in specific situations, such as when the victim is a faceless corporation or an anonymous retail entity rather than a visible person (Pithers, 1999). From the evolutionary game theory and behavioral ecological perspectives, theft is also understood as a highly complex strategy of resource allocation, attempting to acquire the value of a resource while entirely bypassing the cost of obtaining it (Broom and Ruxton, 1998; Barker and Bronstein, 2016; Da Silva and Bonini, 2024; Garbus and Pollack, 2024). The successful execution of such a strategy must rely heavily on social cognition. However, relatively few studies have empirically tested social cognition in criminals, especially in the context of theft recidivism.

A set of questionnaires and psychological tasks could be utilized to address social cognition associated with moral disengagement and contextual empathy deficits that may be related to theft. The Social Norms Questionnaire (SNQ-22) assesses an explicit understanding of and adherence to societal rules (Kramer et al., 2014). In contrast, the Dictator Game (DG) and Dove-Hawk Game (HDG) serve as typical tasks to examine social cognition at resource allocation. Thus, DG assesses prosociality and fairness of a player with discretionary distribution of resources with a passive recipient (Kahneman et al., 1986; Brocklenbank et al., 2011), whereas the HDG is another task to examine social cognition, specifically focusing on conflict resolution, aggression, cooperation, and risk-taking, with competitive resource allocation in which a player chooses between an aggressive, resource-monopolizing strategy (Hawk) and a cooperative, sharing strategy (Dove) against an opponent (Rapoport and Chammah, 1963; Maynard Smith and Price, 1973; Nelissen et al., 2007; Bengart et al., 2021; Lin and Schank, 2022).

The current study aimed to unveil the social cognition deficits present in theft recidivists with kleptomania (TR+K) and instrumental theft recidivists without kleptomania (TR-K), comparing them to control subjects without criminal records (CT), using the SNQ-22 alongside the DG and HDG. Thus, combining these tasks and questionnaire, the social cognition associated with theft recidivism was comprehensively examined by bridging explicit, self-reported moral disengagement with the implicit, behavioral manifestations of contextual empathy deficits in resource-sharing scenarios. We hypothesized that, consistent with the conventional view, theft recidivists exhibited social cognition deficits.

## METHODS

### Subjects

This study was conducted with 53 control participants (CT) who had no history of criminal records and 63 theft recidivists who had prior records of incarceration due to larceny, primarily shoplifting. These participants were recruited for this study, along with our other studies (Goto et al., 2026b; Goto et al., 2026a). TR participants were further divided into those who were diagnosed with kleptomania (TR+K, n=16) and under treatment at the clinic at the time of the investigation and those without the diagnosis (TR-K, n=47). One TR-K participant dropped out from the investigation after obtaining consent but before starting the survey, such that no data was obtained from this participant. Among 16 TR+K and 46 TR-K participants, 3 each of TR+K and TR-K participants accomplished the SNQ-22 but dropped out from the DG and HDG due to loss of interest. The inclusion criteria were ages between 18 and 79 years old and living in Japan at the time of the investigation. The exclusion criteria were the inability to understand the details of the investigation for any reasons, for which none had met. Socioeconomic status was not considered for inclusion/exclusion criteria, as it is relatively invariable among participants, owing to that the TR+K and TR-K participants had been in rehabilitation centers or welfare facilities for their reintegration into society.

### Social Norms Questionnaire (SNQ-22)

A self-report questionnaire (Kramer et al., 2014) was administered to examine how participants recognize social rules in their own cultural backgrounds. Thus, the questionnaire assesses the knowledge of social rules but does not address whether participants follow such social rules. The questionnaire consists of 22 questions, 13 of which are to evaluate whether the participants accept behaviors that are considered socially unacceptable (Break), and 9 of which are to evaluate whether the participants consider that generally acceptable behaviors are wrong (Over-adherence), respectively. Accordingly, the maximum score of the questionnaire is twenty-two (thirteen Break norms score or B-score, and nine Over-adherence score or O-score), and higher the score is, it indicates stronger the recognition of social norms is. The questionnaire was translated into Japanese, along with modification to fit the Japanese cultural backgrounds, and had been used in our previous study (Kaneko et al., 2021). The average score with this Japanese version of the questionnaire administered in healthy Japanese adults was previously shown to comparable to that of the original English version of the questionnaire in US population (Kaneko et al., 2021).

### Dictator Game (DG)

The DG is a socioeconomical psychological task used to assess prosociality and fairness by the discretionary allocation of monetary resources between a participant and an opponent (Kahneman et al., 1986; Brocklenbank et al., 2011). The DG designed in this study began with the presentation of an image (a face photo) to the opponent, followed by trials. The task consisted of six trials with different allocation scenarios. The first three trials were designed to evaluate prosociality (P-phase), in which a choice of allocation was either a split with a larger amount to the opponent than the self (prosocial choice) or an even split between them. The amount of difference in the prosocial choice in each trial was 375, 600, and 50 Japanese yen (approximately corresponding to US$2.50, 4.00, and 0.50), respectively. The later three trials were designed to assess fairness (F-phase), in which a choice of allocation was either an even split (fair choice) or an unfair (selfish) split with larger amount to the self than the opponent. The amount of difference in the unfair choice in each trial was 500, 800, and 50 Japanese yen (approximately corresponding to US$3.00, 5.00, and 0.50), respectively. These amounts of differences in the prosocial and unfair choices were determined based on other studies utilizing DG. The number of prosocial choices in the P-phase and fair choices in the F-phase were measured.

### Hawk-Dove Game (HDG)

The HDG was utilized to assess competitive resource allocation. The HDG is a strategic psychological task frequently used in the context of evolutionary biology (Rapoport and Chammah, 1963; Maynard Smith and Price, 1973), although it can also be used to assess social cognitions, such as social cooperation and fairness (positive reciprocity of social interaction) as well as reiteration and aggression (negative reciprocity of social interaction), given the nature of the task to play with an opponent to compete for the resource (Nelissen et al., 2007; Bengart et al., 2021; Lin and Schank, 2022). The fundamental principle of the HDG is that two people compete for a divisible resource V. Thus, a participant and an opponent (a computer player) choose either to (a) take the resource, that is the Hawk strategy, which fights for the whole resource V at cost C, or (b) wait to see the opponent’s decision, that is the Dove strategy, which concedes the resource to a Hawk or splits the resource with a Dove opponent. Thus, if a participant decides to play Hawk and the opponent plays Dove, the participant can get V and the opponent gets 0, whereas the opponent also plays Hawk, both the participant and opponent experience a devastating cost C (and therefore the payoff is (V-C)/2). In contrast, if a participant decides to play Dove and the opponent plays Hawk, the opponent can get V and the participant gets 0, whereas the opponent also plays Dove, the participant and opponent share the resource (the payoff is V/2). When V > C, Hawk can be a dominant strategy, whereas V < C, neither Hawk nor Dove strategy is stable. In this study, the value of the resource and cost were determined to be C = 2V (e.g., V = 100 Japanese yen and C = 200 Japanese yen). At the beginning of the task, participants were asked to collect as many resources as they could while competing with a computer-generated opponent by choosing either Hawk response (H-response) or Dove response in each trial. The HDG consisted of three phases, with 20 trials in each phase. In each phase, the probability that the computer-generated opponent decided to take the resource varied. The first was the training phase, where the opponent took the Hawk strategy at the probability of 0.5. This phase was aimed at participants getting accustomed to the game. Then, two phases of trials were administered pseudo-randomly, one for trials in which the opponent primarily took the Dove strategy (the probability of Hawk strategy was at 0.2; D-phase), and the other in which the opponent primarily took the Hawk strategy (the probability of Hawk strategy was at 0.8; H-phase). The numbers of Hawk responses (H-response) that participants took against the opponents in D-phase and H-phase were measured.

### Study Design

This study was conducted in accordance with the Declaration of Helsinki and the Ethical Guidelines for Medical and Health Research Involving Human Subjects of the Japanese Ministry of Health, Labour, and Welfare. All procedures were approved by the Human Research Ethics Committee of the Kyoto University Graduate School of Informatics. Written informed consent was obtained from all participants before the investigation. Age, sex, and smoking status were collected at the time of enrollment. In this study, SNQ-22 was first administered, followed by DG and HDG in this sequence.

### Data Analysis

All data are expressed as mean ± standard error of the mean (s.e.m.). Statistical analyses were conducted using JASP ver. 0.97.1 (Team, 2026) and OriginPro ver. 2026 (OriginLab Corporation, Northampton, MA, USA).

Given the non-normal distributions of data from the SNQ-22 and HDG, rank-based inverse normal transformation was applied to convert the non-normal data into standard normal z-scores. A continuous covariate (Age) was also centered and scaled by standardization before statistical analysis.

Data were initially evaluated using frequentist statistical methods. However, when the data for the HDG were analyzed using a Generalized Linear Mixed Model (GLMM), the inclusion of categorical demographic covariates caused model instability and convergence failure, likely due to data sparsity and thin cross-classifications within the smallest cohort (TR+K, n=13). To stabilize parameter estimation and retain these critical covariates, we transitioned to a Bayesian approach, since the Bayesian framework utilizing weakly informative priors provided necessary regularization, preventing unstable estimates caused by sparse subgroups and allowing for robust inference. For consistency in the analysis, Analysis of Covariance (ANCOVA) for the data from the SNQ-22 was also conducted using a Bayesian approach.

#### Bayesian ANCOVA

Bayesian Analysis of Covariance (ANCOVA) was used to test the group effect (Group: TR+K vs. TR-K vs. CT) on the scores of SNQ-22, while controlling for Age (ages of participants), Sex (male vs. female), and Smoking status (yes vs. no), as covariates. To evaluate the evidence, a set of candidate models (Null model, Covariate-only model, Main Effect model, and Main Effect + Covariate model) against the null model was compared. The Jeffreys-Zellner-Siow (JZS) prior, which is a standard default objective prior framework, was used, with a Cauchy prior with a scale of r = 0.5 for the fixed effects and r = 0.354 for the covariates (Rouder and Morey, 2012). Bayes Factors (BF_10_) were evaluated to quantify the likelihood of the data under the alternative model relative to the null model. To interpret the strength of the evidence, the standard classification scheme by Lee and Wagenmakers (Lee and Wagenmakers, 2014) was adopted, where 1 < BF_10_ < 3 indicates anecdotal evidence, 3 < BF_10_ < 10 indicates moderate evidence, and BF_10_ > 10 indicates strong evidence for the alternative hypothesis. In addition, where relevant, the posterior median for effect sizes alongside their 95% credible intervals (CI) was also reported.

#### Bayesian GLMM

Bayesian GLMM was used to handle the hierarchical structure of the data and repeated measures in the data from the DG and HDG. Two separate modeling strategies were employed based on the distribution of the dependent variables. First, to analyze the binary dependent variable (prosocial vs. fair choice in each trial of the P-phase and fair vs. unfair choice in each trial of the F-phase) in the DG, Bayesian GLMM utilizing a Bernoulli distribution with a logit link function was adopted, whereas for the HDG to analyze the continuous, normally distributed dependent variable (the number of H-response), Bayesian GLMM utilizing a Gaussian distribution with an identity link function was specified. Both models included Group (TR+K vs. TR-K vs. CT), Age (ages of participants), Sex (male vs. female), Smoking (yes vs. no), and Phase (DG: trial 1 to 3 in either P-phase or F-phase; HDG: D-phase vs. H-phase), as fixed effects. To account for within-subject dependencies, a random intercept for SubjectID (each participant) was included. We utilized prior distributions set as a default by the Bayesian GLMM module in JASP, which are designed to be weakly informative and provide regularization to prevent overfitting while allowing the data to drive the posterior estimates. Posterior distributions were estimated using Markov Chain Monte Carlo (MCMC) sampling. For each model, the sampler was run across three independent chains with 4,000 iterations per chain. The first 2,000 iterations were discarded as burn-in (warmup). Model convergence and sampling quality were verified using the Gelman-Rubin diagnostic (*R̂* < 1.05 for all parameters) and Effective Sample Size (ESS bulk and tail > 400) for all parameters, which indicates that the independent chains successfully converged to the same target distribution. Posterior distributions were summarized using the posterior mean (Estimate) and 95% Credible Intervals (95% CI). Pairwise comparisons were evaluated using custom contrast weights. A fixed effect was considered statistically credible if its 95% CI excluded zero.

## RESULTS

### Social Norms

SNQ-22 was administered to evaluate the recognition of social norms (Fig. 1a; Suppl. Table S1). The questionnaire comprised of a Break norms score (B-score), indicating that the subject accepts behaviors that are socially unacceptable, and an Over-adherence score (O-score), indicating that the subject considers that generally acceptable behaviors are wrong.

**Figure 1.**
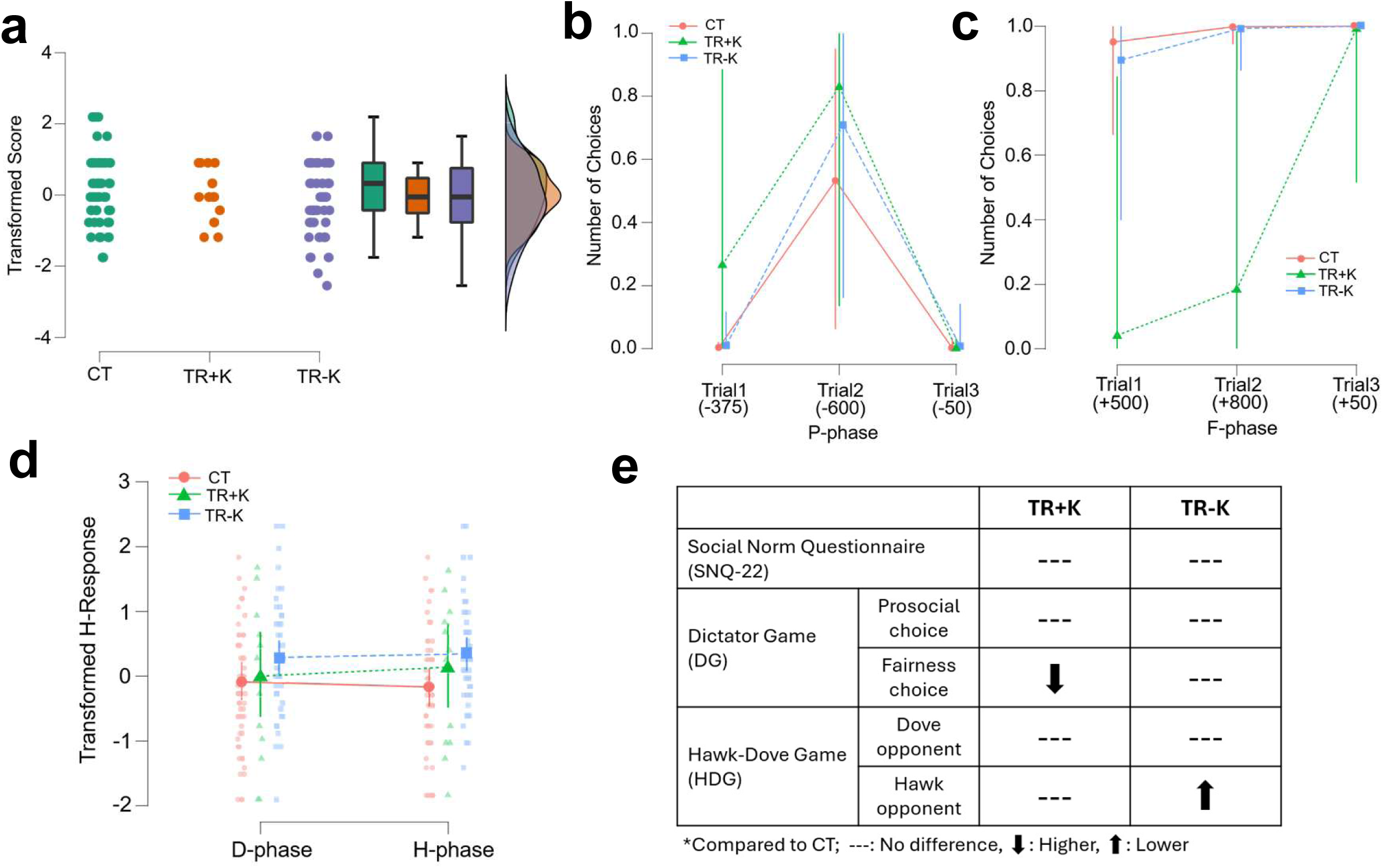
Comparisons of TR+K, TR-K, and CT participants in SNQ-22, DG, and HDG. **a.** A raincloud plot illustrating the total SNQ-22 scores with the rank-based inverse normal transformation for TR+K, TR-K, and CT. **b, c.** Graphs showing the number of prosocial choices in the P-phase (b) and fair choices in the F-phase (c), respectively, of DG. Parentheses indicate the amount of allocation difference (relative to the opponent, so that a minus indicates larger allocation to the opponent than self, and vice versa) in Japanese yen. Error bars indicate 95% CI. **d.** A graph showing the number of H-responses with the rank-based inverse normal transformation in D-phase (the opponent with the dove strategy) and H-phase (the opponent with the hawk strategy) of HDG. The dark circle, triangle, square, and error bars indicate the means and 95% CI for each of CT, TR+K, and TR-K, respectively. Light-color circles, triangles, and squares indicate the H-responses of each participant. **e.** A summary illustrating how the measurements of SNQ-22, DG, and HDG in TR+K and TR-K differ from those of CT.

Bayesian ANCOVA was conducted to determine the effect of Group (TR+K vs. TR-K vs. CT) on the total scores (combining B- and O-scores) while controlling for Age, Sex, and Smoking (Suppl. Table S2). The analysis revealed that the best-fitting model included only Sex (BF_10_ = 19.19). The analysis of effects yielded strong evidence for Sex (BF_incl_ = 11.41), but not for other predictors, including Group (BF_incl_ = 0.146). Strong evidence of a credible effect was observed for stricter adherence in female than male participants (female posterior mean = 0.246, 95% CI [0.062, 0.429]; Suppl. Table S2) in single model inference. Bayesian ANCOVA was also conducted for B-score (Suppl. Table S3) and O-score (Suppl. Table S4), respectively. The analysis revealed that no model was higher than BF_10_ of 1.0 for B-score, suggesting that the observed data are less likely under the alternative than the null hypothesis, whereas the best-fitting model for O-score included Sex (BF_10_ = 3.976). Moderate evidence of effect was observed for Sex (BF_incl_ = 3.637) in the analysis of effects. Strong evidence of a credible effect was also observed for stricter adherence in female than male participants (female posterior mean = 0.225, 95% CI [0.045, 0.423]; Suppl. Table S4) in single model inference.

These results suggest that female participants, regardless of theft recidivists or control participants, recognized social norms that were stricter than those of male participants. No difference in the recognition of social norms between groups suggest a dissociation between explicit knowledge and the behavioral manifestation of social norms in theft recidivists.

### Discretionary Resource Allocation

The DG was administered to assess discretionary resource allocation. The game consisted of two phases, one for prosocial trials (P-phase; Fig. 1b, Suppl. Table S5), where a participant chose either an even split allocation with the opponent or a split with a larger amount to the opponent than self (prosocial choice), and the other for fair trials (F-phase; Fig. 1c, Suppl. Table S5), where the participant chose either an even split allocation with the opponent (fair choice) or a split with a larger amount to self than the opponent. Given that prosociality and fairness have been shown to be distinct processes (Leiberg et al., 2011; Kim et al., 2023), supported by that there was no correlation between the number of prosocial choices in P-phase and fair choices in F-phase (Kendall’s correlation coefficient τ = -0.117, p = 0.170), P-phase and F-phase were separately analyzed.

Bayesian logit-link GLMM (Bernoulli family) was conducted to examine the group effect on the number of prosocial and fair choices in the P-phase and F-phase, respectively. Fixed effects comprised Group (TR+K vs. TR-K vs. CT), Age, Sex, Smoking, Phase (3 trials each for P-phase or F-phase), and the Group x Phase interaction term, while SubjectID (each participant) was modeled as a random intercept. MCMC sampling achieved robust convergence across all model parameters (*R̂* ≤ 1.005, effective sample size (ESS) ≥ 494.9). Age did not exert a credible effect on the outcome (Estimate = 0.014, 95% CI [-0.073, 0.102] in P-phase; Estimate = 0.030, 95% CI [-0.021, 0.086] in F-phase). Similarly, neither sex (Female: Estimate = 0.472, 95% CI [-3.887, 4.822] in P-phase; Estimate = 0.903, 95% CI [-1.589, 3.544] in F-phase; Male: Estimate = 0.857, 95% CI [-3.701, 5.595] in P-phase; Estimate = 1.948, 95% CI [-0.831, 4.991] in F-phase) nor smoking status (Non-smoker: Estimate = 0.533, 95% CI [-4.489, 5.400] in P-phase; Estimate = 1.904, 95% CI [-0.897, 4.960] in F-phase; Smoker: Estimate = 0.796, 95% CI [-3.046, 4.914] in P-phase; Estimate = 0.947, 95% CI [-1.614, 3.642] in F-phase) showed credible posterior differences from zero.

In P-phase (Suppl. Table S6, Fig. 1b), the overall main effects for Group (CT: Estimate = -0.611, 95% CI [-5.147, 3.675]; TR+K: Estimate = 1.415, 95% CI [-3.177, 6.747]; TR-K: Estimate = 1.189, 95% CI [-3.881, 6.149]) spanned zero. However, group performance varied across phases. Thus, evaluation of the Group x Phase interaction term revealed that both the TR+K (Estimate = 5.911, 95% CI [0.520, 13.000]) and TR-K (Estimate = 5.172, 95% CI [0.054, 11.400]) exhibited credible positive parameter estimates, whereas CT remained uncredible (Estimate = 4.289, 95% CI [-0.204, 9.725]) in the trial where the largest amount was allocated to the opponent by a prosocial choice (Fig. 1b). However, direct pairwise contrasts revealed no credible differences between CT, TR+K, and TR-K in any trials (all 95% highest probability distribution (HPD) intervals spanned zero). Together, these results indicate that while the baseline response was robust across the cohort, neither TR+K nor TR-K was different from CT in prosocial choices.

In F-phase (Suppl. Table S7, Fig. 1c), evaluation of Group relative to the model intercept indicated no credible effects for CT (Estimate = 4.814, 95% CI [-0.793, 11.07]), TR+K (Estimate = -1.100, 95% CI [-7.456, 5.308]), and TR-K (Estimate = 4.132, 95% CI [-1.433, 10.59]). Evaluation of the Group x Phase interaction term revealed that both the CT (Estimate = 7.701, 95% CI [1.398, 15.31]) and TR-K (Estimate = 7.569, 95% CI [1.047, 15.52]) exhibited credibly positive parameter estimates, whereas TR+K remained uncredible (Estimate = 3.924, 95% CI [-3.415, 12.22]) in the trial where the smallest amount was allocated to the self by an unfair choice. Pairwise contrasts confirmed credibly negative effects between TR+K and CT in specific trials (Estimate = -0.840, 95% HPD [-1.000, -0.081] in trial 1; Estimate = -0.795, 95% HPD [-1.000, -0.010] in trial 2), whereas no credible overall difference was observed between CT to TR-K, suggesting that TR+K resulted in a credibly distinct outcome compared to the CT.

Collectively, these results suggest that the prosociality of TR+K and TR-K is not different from that of CT, whereas TR-KA, but not TR-NK, exhibits decreased fairness compared to CT at discretionary resource allocation.

### Competitive Resource Allocation

The HDG was administered to assess competitive resource allocation (Suppl. Table S8). Two phases of the HDG, one for trials in which participants competed with the Dove (cooperative/equity) strategy opponent for the resources (D-phase), and the other for trials in which participants competed with the Hawk (aggressive/inequity) strategy opponent for the resources (H-phase), were subjected to analysis.

Bayesian Identity-link GLMM (Gaussian family) was utilized to examine the effects of Group, Phase (D-phase vs. H-phase), and Group x Phase interaction on the number of Hawk responses that participants made (H-response), while adjusting for Age, Sex, and Smoking, and SubjectID was modeled as a random intercept (Suppl. Table S9, Fig. 1d). Model diagnostics indicated robust chain convergence (*R̂* ≤ 1.003) and adequate effective sample sizes (ESS ≥ 1244). None of the main effects of the covariates, such as Age, Sex, or Smoking, demonstrated a credible effect on the outcome, as all 95% credible intervals spanned zero. Analysis of the estimated marginal means revealed that the TR-K maintained a credibly positive effect under both opponent phases (Median = 0.289, 95% HPD [0.028, 0.549] in D-phase; Median = 0.343, 95% HPD [0.079, 0.602] in H-phase). In addition, contrast analyses indicated a credible effect where TR-K was significantly different from CT in H-phase (Estimate = 0.508, 95% HPD [0.118, 0.915]). Insufficient credibility was observed for differences between the other groups. These observations were further supported by the results of GLMM analysis for payoffs for participants in the HDG, indicating that the payoffs of TR-K in the H-phase are lower than those of CT (Estimate = -0.550, 95% HPD [-0.966, -0.158]; Suppl. Table S10).

Collectively, these results suggest that TR-K is inflexible in their response strategy, demonstrating more aggression even in high-cost competition, at resource allocation.

## DISCUSSION

The current study revealed distinct social cognition profiles among the participant groups (Fig. 1e). Thus, there were no differences in recognition of social rules as assessed by the SNQ-22 between groups, suggesting that morality and rule comprehension of TR+K and TR-K may be comparable to those of CT. In contrast, at the discretionary resource allocation with others tested in the DG, while all groups displayed comparable prosocial choices, TR+K made fewer fair choices than CT and TR-NK. Moreover, in the competitive resource allocation with others examined in the HDG, TR-K demonstrated more aggressive responses to the aggressive (Hawk strategy) opponent than TR+K and CT, while no differences emerged between the groups against cooperative (Dove strategy) opponent. These observations suggest that recurrent theft may not be caused by a generalized lack of moral knowledge, but the “knowing vs. doing” gap (Pfeffer and Sutton, 1999) may play a significant role.

Discrepant performances of TR+K and TR-K in the DG and HDG suggest that distinct mechanisms may affect their social cognition. Such distinct mechanisms appear to be in line with our recent findings that both TR+K and TR-K exhibit higher impulsivity than CT (Goto et al., 2026b), whereas TR+K, but not TR-K, demonstrates heightened loss aversion (Goto et al., 2026a), and can be explained by the mechanical interaction of such heightened impulsivity and loss aversion. Thus, at discretionary resource allocation in the DG, the participants were handed a pool of resources and asked to divide them. Consequently, for individuals with heightened loss aversion, such as TR+K, a resource allocation is psychologically processed as a painful loss rather than a neutral division, making them choose fewer fair choices in the DG. Nonetheless, no credible evidence of a difference in the number of prosocial choices was observed between the groups. This could be explained by distinct processes of prosociality and fairness in the dual process model, with prosociality in a hot, emotional process vs. fairness in a cold, cognitive process (Leiberg et al., 2011; Kim et al., 2023). In fact, although no credible evidence was found, percentages of prosocial choices in the DG appear to be higher in TR+K and TR-K than in CT (Suppl. Table S5). Such an observation may be explained by the facilitation of emotional prosocial reactions with higher impulsivity. Conversely, the HDG is a game that stimulates interpersonal social conflict and threat (risk of devastating loss) for resource competition. In such a competitive context, higher impulsivity instinctively drives to seize immediate resource acquisition via the aggressive Hawk strategy. In particular, such aggression could be augmented when faced with an adversarial aggressive opponent as righteous retaliation, causing TR-K to make more aggressive responses to the aggressive opponent in the HDG. In contrast, heightened loss aversion may act as an emergency brake for such aggression, as the threat of a devastating Hawk-Hawk penalty is highly salient and intense fear of severe loss, forcing TR+K to play cautiously and successfully matching the strategic behavior of normal subjects. Collectively, these behavioral profiles explain that “internal” threat causing reflex to protect themselves from feeling deprived may play a role in TR+K, whereas “external” threat causing necessary defense against perceived hostility may be in TR-K.

These findings suggest that moral disengagement and contextual empathy deficits may ultimately operate through distinct mechanisms in theft recidivists. A contextual empathy deficit is that a person can feel empathy normally, but specific environments or triggers cause the empathy to temporarily shut down (Pithers, 1999). Thus, in TR+K, such a deficit may involve an internal threat triggered by the prospect of losing a resource, which temporarily blinds their otherwise intact empathy, whereas in TR-K, the empathy switch turns off in the presence of an external threat or interpersonal conflict. Moral disengagement is the cognitive rationalization process of convincing oneself that ethical standards do not apply in a specific situation (Bandura, 1991). TR+K, driven by an internal threat with heightened loss aversion and impulsivity, likely employ mechanisms like distortion of consequences (“*The store makes millions, they won’t miss this one item*”) or advantageous comparison (“*I’m not hurting anyone physically, I’m just taking this*”). This may also explain the ego-dystonic nature of kleptomania, where individuals feel intense guilt because their moral engagement was intact, but they lacked the brakes to stop the impulse. In contrast, TR-K, driven by an external threat with impulsivity, likely use moral disengagement mechanisms, such as displacement of responsibility or blaming the victim (“*Society is rigged against me*,” or “*Since the salesclerks are rude to me, they deserve what they get*”).

This study has several major limitations. First, the sample size, particularly for TR+K, was relatively small, necessitating the use of a Bayesian analytical approach to ensure robust parameter estimation. Second, the cross-sectional design of the study makes causal relationships between social cognition deficits and theft recidivism unclear. In addition, the neuronal mechanisms that mediate these social behaviors are not fully elucidated in the current study. The prefrontal cortex (PFC) plays an important role in the neurobiological mechanisms of social cognition (Forbes and Grafman, 2010). The PFC mediates executive functioning (Friedman and Robbins, 2022), impulse control (Kim and Lee, 2011), moral reasoning (Forbes and Grafman, 2010), and the integration of social norms into decision-making (Christian and Soutschek, 2022). Functional neuroimaging studies demonstrate that the PFC is involved in social cognition relevant to the DG and HDG. In particular, regions such as the dorsolateral and ventromedial PFC exhibit increased activation when individuals suppress selfish urges to make fair divisions (Baumgartner et al., 2011) or when they successfully navigate the cooperative demands of conflict resolution (Pretus et al., 2019). Our previous studies suggest that an imbalance in left and right prefrontal cortex (PFC) activity may be associated with heightened loss aversion in kleptomania (Goto et al., 2026a), whereas altered medial PFC activity may be related to impulsivity in theft recidivists (Goto et al., 2026b). However, the exact mechanisms by which these brain areas mediate the observed behavioral discrepancies between TR+K and TR-K require further investigation.

In conclusion, this study demonstrates that deficits in moral knowledge or general prosocial capabilities may not be primary causes of offenses in theft recidivists. Instead, recidivism may stem from contextual failures associated with social cognition deficits affected by the interplay of impulsivity and loss aversion, which are distinctly in kleptomanic and instrumental theft recidivists. Specifically, recurrent theft with kleptomania may be rooted in heightened loss aversion, which hijack decision-making during resource allocation, whereas instrumental theft recidivism without kleptomania may be associated with impulsivity overriding risk in conflict. Thus, these findings suggest the importance of targeted, context-specific rehabilitation and therapeutic interventions rather than generalized punitive measures for preventing recidivism.

## Funding

This work was supported by the Japan Society for the Promotion of Science (JSPS) Grant-in-Aid for Scientific Research (B) 25K00898, awarded to YG.

## Data Availability

The data underlying this article will be shared upon reasonable request from the corresponding author.

## Author Contributions

YG contributed to the conceptualization, methodology, formal analysis, investigation, data curation, project administration, writing, visualization, validation, supervision, and funding acquisition. MIC contributed to the formal analysis, writing, and visualization. SA contributed to the conceptualization, resources, and writing. CK contributed to the conceptualization, resources, and writing. MJW contributed to the conceptualization, resources, and writing. YAL contributed to the formal analysis and writing. All authors have reviewed, edited, and approved the manuscript prior to its submission.

## Competing Interests

The authors declare no conflict of interest.

## Acknowledgements

We would like to thank the staff of the Non-Profit Organization Kurashi O-en Network (Nagoya, Japan), Nishi Hongwanji Byakkoso (Kyoto, Japan), Non-Profit Organization Kyoto MAC (Kyoto, Japan), Kyoto Hogo Ikusei Kai (Kyoto, Japan), and Liberty Women’s House Olive (Otsu, Japan) for recruiting participants, scheduling surveys, and providing various technical supports.

## Notes

### Competing Interest Statement

The authors have declared no competing interest.

