## Supplementary Materials for "Altered Social Cognition Associated with Kleptomanic and Instrumental Thefts"

Altered Social Cognition in Theft Recidivists with and without Kleptomania

**Supplementary Table S1. A summary of SNQ-22 scores.**

**Supplementary Table S2. Bayesian ANCOVA for total scores in SNQ-22.**

**Supplementary Table S3. Bayesian ANCOVA for B-scores in SNQ-22.**

**Supplementary Table S4. Bayesian ANCOVA for O-scores in SNQ-22.**

**Supplementary Table S5. A summary of measurements in DG.**

**Supplementary Table S6. Bayesian GLMM for prosocial choice in P-phase of DG.**

**Supplementary Table S7. Bayesian GLMM for the fair choice in F-phase of DG.**

**Supplementary Table S8. A summary of measurements in HDG.**

**Supplementary Table S9. Bayesian GLMM for H-response in HDG.**

**Supplementary Table S10. Bayesian GLMM for payoffs in HDG.**

**Supplementary Table S1. A summary of SNQ-22 scores.**

| | CT | Theft Recidivist | |
|---|---|---|---|
| | | TR+K | TR-K |
| **Total Score** | 17.21 ± 0.29 | 16.94 ± 0.43 | 16.52 ± 0.35 |
| **B-score** | 11.91 ± 0.16 | 11.88 ± 0.24 | 11.80 ± 0.21 |
| **O-score** | 5.30 ± 0.33 | 5.06 ± 0.44 | 4.72 ± 0.36 |

SNQ-22 – Social norms questionnaire; CT – Control subject with no criminal record; TR+K – Theft recidivist with kleptomania; TR-K – Theft recidivist without kleptomania; B-score – Break norm score; O-score – Over-adherence norm score; Total score – B- and O-scores combined; mean ± s.e.m.

### Supplementary Table S2. Bayesian ANCOVA for total scores in SNQ-22.

**Model Comparison (Showing the models with $BF_{10} \geq 1.0$)**

| Models | P(M) | P(M\|data) | $BF_M$ | $BF_{10}$ | error% |
|---|---|---|---|---|---|
| Null model | 0.063 | 0.025 | 0.382 | 1.000 | |
| Sex | 0.063 | 0.477 | 13.66 | 19.19 | $1.622 \times 10^{-7}$ |
| Sex + Smoking | 0.063 | 0.145 | 2.546 | 5.842 | 0.863 |
| Sex + Age | 0.063 | 0.143 | 2.505 | 5.761 | 0.847 |
| Sex + Group | 0.063 | 0.071 | 1.141 | 2.845 | 1.251 |
| Sex + Smoking + Age | 0.063 | 0.042 | 0.651 | 1.675 | 1.228 |
| Smoking | 0.063 | 0.030 | 0.464 | 1.208 | 0.012 |

**Analysis of Effects**

| Effects | P(incl) | P(excl) | P(incl\|data) | P(excl\|data) | $BF_{incl}$ |
|---|---|---|---|---|---|
| Group | 0.500 | 0.500 | 0.126 | 0.874 | 0.145 |
| Sex | 0.500 | 0.500 | 0.920 | 0.080 | 11.46 |
| Smoking | 0.500 | 0.500 | 0.251 | 0.749 | 0.335 |
| Age | 0.500 | 0.500 | 0.225 | 0.775 | 0.290 |

**Single Model Inference**

| Variable | Level | Mean | SD | 95% CI | |
|---|---|---|---|---|---|
| | | | | Lower | Upper |
| Intercept | | -0.087 | 0.101 | -0.300 | 0.092 |
| Group | CT | 0.085 | 0.117 | -0.141 | 0.312 |
| | TR+K | -0.034 | 0.143 | -0.314 | 0.242 |
| | TR-K | -0.052 | 0.119 | -0.284 | 0.179 |
| Sex | Female | 0.242 | 0.097 | 0.060 | 0.435 |
| | Male | -0.242 | 0.097 | -0.435 | -0.060 |
| Smoking | No | 0.034 | 0.103 | -0.161 | 0.243 |
| | Yes | -0.034 | 0.103 | -0.243 | 0.161 |
| Age | | -0.004 | 0.005 | -0.015 | 0.006 |

P(M) – Prior model probability; P(M|data) – Posterior model probability; $BF_M$ – Model Bayes factor; $BF_{10}$ – Bayes factor ($H_1$ over $H_0$); Error% – Numerical error percentage; P(incl) – Prior inclusion probability; P(excl) – Prior exclusion probability; P(incl|data) – Posterior inclusion probability; P(excl|data) – Posterior exclusion probability; $BF_{incl}$ – Inclusion Bayes factor; SD – Standard deviation; CI – Credible interval

### Supplementary Table S3. Bayesian ANCOVA for B-scores in SNQ-22.

**Model Comparison (Showing the models with $BF_{10} \geq 1.0$)**

| Models | P(M) | P(M\|data) | $BF_M$ | $BF_{10}$ | error% |
|---|---|---|---|---|---|
| Null model | 0.063 | 0.389 | 9.543 | 1.000 | |

**Analysis of Effects**

| Effects | P(incl) | P(excl) | P(incl\|data) | P(excl\|data) | $BF_{incl}$ |
|---|---|---|---|---|---|
| Group | 0.500 | 0.500 | 0.091 | 0.909 | 0.100 |
| Sex | 0.500 | 0.500 | 0.178 | 0.822 | 0.216 |
| Smoking | 0.500 | 0.500 | 0.326 | 0.674 | 0.485 |
| Age | 0.500 | 0.500 | 0.230 | 0.770 | 0.298 |

**Single Model Inference**

| Variable | Level | Mean | SD | 95% CI | |
|---|---|---|---|---|---|
| | | | | Lower | Upper |
| Intercept | | -0.080 | 0.095 | -0.270 | 0.094 |
| Group | CT | 0.002 | 0.112 | -0.210 | 0.233 |
| | TR+K | -0.045 | 0.134 | -0.320 | 0.209 |
| | TR-K | 0.043 | 0.111 | -0.178 | 0.256 |
| Sex | Female | -0.016 | 0.087 | -0.184 | 0.156 |
| | Male | 0.016 | 0.087 | -0.156 | 0.184 |
| Smoking | No | 0.117 | 0.095 | -0.059 | 0.309 |
| | Yes | -0.117 | 0.095 | -0.309 | 0.059 |
| Age | | -0.004 | 0.005 | -0.013 | 0.005 |

### Supplementary Table S4. Bayesian ANCOVA for O-scores in SNQ-22.

**Model Comparison (Showing the models with $BF_{10} \geq 1.0$)**

| Models | P(M) | P(M\|data) | $BF_M$ | $BF_{10}$ | error% |
|---|---|---|---|---|---|
| Null model | 0.063 | 0.016 | 1.960 | 1.000 | |
| Sex | 0.063 | 0.460 | 12.75 | 3.976 | 0.006 |

**Analysis of Effects**

| Effects | P(incl) | P(excl) | P(incl\|data) | P(excl\|data) | $BF_{incl}$ |
|---|---|---|---|---|---|
| Group | 0.500 | 0.500 | 0.128 | 0.872 | 0.146 |
| Sex | 0.500 | 0.500 | 0.783 | 0.217 | 3.607 |
| Smoking | 0.500 | 0.500 | 0.196 | 0.804 | 0.243 |
| Age | 0.500 | 0.500 | 0.178 | 0.822 | 0.217 |

**Single Model Inference**

| Variable | Level | Mean | SD | 95% CI | |
|---|---|---|---|---|---|
| | | | | Lower | Upper |
| Intercept | | -0.062 | 0.099 | -0.259 | 0.136 |
| Group | CT | 0.094 | 0.117 | -0.114 | 0.340 |
| | TR+K | -0.039 | 0.147 | -0.341 | 0.272 |
| | TR-K | -0.055 | 0.119 | -0.287 | 0.272 |
| Sex | Female | 0.225 | 0.099 | 0.045 | 0.423 |
| | Male | -0.225 | 0.099 | -0.423 | -0.045 |
| Smoking | No | -0.037 | 0.101 | -0.248 | 0.162 |
| | Yes | 0.037 | 0.101 | -0.162 | 0.248 |
| Age | | -0.003 | 0.005 | -0.013 | 0.007 |

### Supplementary Table S5. A summary of measurements in DG.

| Measurement (Phase) | Trial | CT | Theft Recidivist | |
|---|---|---|---|---|
| | | | TR+K | TR-K |
| **Prosocial Choice (P-phase)** | **Trial 1 (-375)** | 5.66% | 38.46% | 23.26% |
| | **Trial 2 (-600)** | 49.06% | 61.54% | 53.49% |
| | **Trial 3 (-50)** | 15.09% | 7.69% | 27.91% |
| **Fair Choice (F-phase)** | **Trial 1 (+500)** | 69.81% | 30.77% | 62.79% |
| | **Trial 2 (+800)** | 77.36% | 38.46% | 72.09% |
| | **Trial 3 (+50)** | 88.68% | 69.23% | 86.05% |

Percentages of participants who chose a prosocial over a fair allocation choice in P-phase and a fair over an unfair allocation choice in F-phase, respectively. Parentheses indicate the amount of allocation difference in Japanese Yen (relative to the opponent) consequent to choosing either a prosocial (in P-phase) or unfair (in F-phase) choice. DG – Dictator game; CT – Control subject with no criminal record; TR+K – Theft recidivist with kleptomania; TR-K – Theft recidivist without kleptomania.

### Supplementary Table S6. Bayesian GLMM for prosocial choice in P-phase of DG.

**Posterior Estimate Summaries for Fixed Effects and Interaction Term (Difference from intercept)**

| Variable | Level 1 | Level 2 | Estimate | Est.Error | 95% CI | | $\hat{R}$ | ESS(bulk) | ESS(tail) |
|---|---|---|---|---|---|---|---|---|---|
| | | | | | Lower | Upper | | | |
| Intercept | | | -4.154 | 2.340 | -9.317 | 0.308 | 1.000 | 1452 | 2817 |
| Group | CT | | -0.611 | 2.147 | -5.147 | 3.675 | 1.000 | 2092 | 2849 |
| | TR+K | | 1.415 | 2.259 | -3.177 | 6.747 | 1.001 | 2280 | 2442 |
| | TR-K | | 1.189 | 2.298 | -3.881 | 6.149 | 1.002 | 1278 | 2352 |
| Age | | | 0.014 | 0.040 | -0.073 | 0.102 | 1.001 | 1673 | 2541 |
| Sex | Female | | 0.472 | 2.047 | -3.887 | 4.822 | 1.002 | 1118 | 2805 |
| | Male | | 0.857 | 2.051 | -3.701 | 5.595 | 1.000 | 2284 | 2349 |
| Smoking | No | | 0.533 | 2.273 | -4.489 | 5.400 | 1.001 | 1645 | 2532 |
| | Yes | | 0.796 | 1.866 | -3.046 | 4.914 | 1.000 | 1978 | 3054 |
| Phase | Trial 1 | | -0.247 | 2.364 | -5.290 | 4.405 | 1.003 | 851.1 | 2337 |
| | Trial 2 | | 5.124 | 2.359 | 0.823 | 10.49 | 1.001 | 1479 | 2907 |
| | Trial 3 | | -2.884 | 2.256 | -7.999 | 1.552 | 1.003 | 1881 | 2572 |
| Group x Phase | CT | Trial 1 | -3.078 | 2.948 | -9.962 | 2.385 | 1.004 | 794.3 | 2085 |
| | CT | Trial 2 | 4.289 | 2.383 | -0.204 | 9.725 | 1.000 | 1818 | 3342 |
| | CT | Trial 3 | -3.043 | 2.702 | -9.125 | 1.761 | 1.003 | 1766 | 2881 |
| | TR+K | Trial 1 | 3.051 | 2.666 | -2.361 | 9.131 | 1.001 | 1865 | 2611 |
| | TR+K | Trial 2 | 5.911 | 3.007 | 0.520 | 13.00 | 1.001 | 1800 | 2656 |
| | TR+K | Trial 3 | -4.716 | 3.612 | -12.71 | 2.021 | 1.002 | 2187 | 2903 |
| | TR-K | Trial 1 | -0.713 | 2.672 | -6.949 | 4.734 | 1.004 | 789.3 | 1977 |
| | TR-K | Trial 2 | 5.172 | 2.708 | 0.054 | 11.40 | 1.001 | 1732 | 2857 |
| | TR-K | Trial 3 | -0.892 | 2.599 | -6.710 | 4.412 | 1.001 | 1835 | 2526 |

**Estimated marginal means**

| Group | Phase | Median | 95% HPD | |
|---|---|---|---|---|
| | | | Lower | Upper |
| CT | Trial 1 | $9.747 \times 10^{-4}$ | $2.491 \times 10^{-9}$ | 0.022 |
| CT | Trial 2 | 0.533 | 0.062 | 0.951 |
| CT | Trial 3 | 0.001 | $1.289 \times 10^{-9}$ | 0.033 |
| TR+K | Trial 1 | 0.266 | $3.809 \times 10^{-6}$ | 0.887 |
| TR+K | Trial 2 | 0.831 | 0.135 | 1.000 |
| TR+K | Trial 3 | $2.155 \times 10^{-4}$ | $6.620 \times 10^{-13}$ | 0.033 |
| TR-K | Trial 1 | 0.010 | $1.143 \times 10^{-6}$ | 0.117 |
| TR-K | Trial 2 | 0.709 | 0.160 | 1.000 |
| TR-K | Trial 3 | 0.008 | $2.154 \times 10^{-8}$ | 0.142 |

**Pairwise Contrasts**

| Contrasts | Estimate | 95% HPD | |
|---|---|---|---|
| | | Lower | Upper |
| CT vs. TR+K, Trial 1 | 0.261 | -0.004 | 0.891 |
| CT vs. TR+K, Trial 1 | 0.007 | -0.028 | 0.119 |
| CT vs. TR+K, Trial 2 | 0.209 | -0.444 | 0.927 |
| CT vs. TR-K, Trial 2 | 0.138 | -0.560 | 0.857 |
| CT vs. TR-A, Trial 3 | $-2.412 \times 10^{-4}$ | -0.048 | 0.045 |
| CT vs. TR-K, Trial 3 | 0.005 | -0.049 | 0.154 |

Est.Error – Estimate error (Posterior standard deviation); $\hat{R}$ – Gelman-Rubin convergence diagnostic; CI – Credible interval; HPD – Highest probability distribution; ESS – Effective sample size; Bulk – Sampling efficiency in the central region of the posterior distribution; Tail – Sampling efficiency in the outer edges of the posterior distribution; The allocation differences (relative to the opponent) are -375, -600, -50 Japanese yen in Trial 1-3, respectively.

### Supplementary Table S7. Bayesian GLMM for the fair choice in F-phase of DG.

**Posterior Estimate Summaries for Fixed Effects and Interaction Term (Difference from intercept)**

| Variable | Level 1 | Level 2 | Estimate | Est.Error | 95% CI | | $\hat{R}$ | ESS(bulk) | ESS(tail) |
|---|---|---|---|---|---|---|---|---|---|
| | | | | | Lower | Upper | | | |
| Intercept | | | 1.271 | 2.694 | -4.206 | 7.479 | 1.003 | 1524 | 498.6 |
| Group | CT | | 4.814 | 2.869 | -0.793 | 11.07 | 1.002 | 1540 | 494.9 |
| | TR+K | | -1.100 | 3.112 | -7.456 | 5.308 | 1.001 | 2207 | 2660 |
| | TR-K | | 4.132 | 2.984 | -1.433 | 10.59 | 1.001 | 2063 | 2407 |
| Age | | | 0.054 | 0.052 | -0.048 | 0.168 | 1.002 | 1835 | 2485 |
| Sex | Female | | 1.785 | 2.547 | -3.791 | 7.253 | 1.004 | 1494 | 500.7 |
| | Male | | 3.445 | 2.822 | -2.188 | 9.772 | 1.002 | 2042 | 2402 |
| Smoking | No | | 3.320 | 2.805 | -2.194 | 9.524 | 1.001 | 2238 | 2626 |
| | Yes | | 1.910 | 2.583 | -3.948 | 7.479 | 1.003 | 1416 | 510.6 |
| Trial | Trial 1 | | -0.580 | 2.550 | -5.900 | 4.682 | 1.003 | 1877 | 522.7 |
| | Trial 2 | | 2.028 | 2.694 | -3.162 | 7.646 | 1.004 | 1932 | 2787 |
| | Trial 3 | | 6.398 | 3.027 | 0.787 | 13.06 | 1.001 | 2012 | 2441 |
| Group x Phase | CT | Trial1 | 1.793 | 2.89 | -4.413 | 7.854 | 1.005 | 1274 | 506.9 |
| | | Trial 2 | 4.949 | 3.196 | -0.903 | 11.91 | 1.001 | 2087 | 3078 |
| | | Trial 3 | 7.701 | 3.478 | 1.398 | 15.31 | 1.002 | 1698 | 551.5 |
| | TR+K | Trial1 | -4.49 | 3.505 | -12.23 | 2.647 | 1.001 | 2172 | 3214 |
| | | Trial 2 | -2.736 | 3.739 | -10.41 | 4.811 | 1.000 | 2559 | 2946 |
| | | Trial 3 | 3.924 | 3.869 | -3.415 | 12.22 | 1.000 | 2419 | 2641 |
| | TR-K | Trial1 | 0.958 | 2.943 | -4.921 | 7.021 | 1.000 | 2340 | 2967 |
| | | Trial 2 | 3.869 | 3.291 | -2.161 | 11.04 | 1.002 | 1878 | 2470 |
| | | Trial 3 | 7.569 | 3.529 | 1.047 | 15.52 | 1.000 | 2250 | 2478 |

**Estimated marginal means**

| Group | Phase | Median | 95% HPD | |
|---|---|---|---|---|
| | | | Lower | Upper |
| CT | Trial 1 | 0.951 | 0.662 | 1.000 |
| CT | Trial 2 | 0.998 | 0.944 | 1.000 |
| CT | Trial 3 | 1.000 | 0.994 | 1.000 |
| TR+K | Trial1 | 0.042 | $4.756 \times 10^{-8}$ | 0.844 |
| TR+K | Trial 2 | 0.184 | $6.139 \times 10^{-7}$ | 0.984 |
| TR+K | Trial 3 | 0.993 | 0.515 | 1.000 |
| TR-K | Trial1 | 0.895 | 0.399 | 1.000 |
| TR-K | Trial 2 | 0.993 | 0.862 | 1.000 |
| TR-K | Trial 3 | 1.000 | 0.993 | 1.000 |

**Pairwise Contrasts**

| Contrasts | Estimate | 95% HPD | |
|---|---|---|---|
| | | Lower | Upper |
| CT vs. TR+K, Trial 1 | -0.840 | -1.000 | -0.081 |
| CT vs. TR+K, Trial 1 | -0.038 | -0.605 | 0.364 |
| CT vs. TR+K, Trial 2 | -0.795 | -1.000 | -0.010 |
| CT vs. TR-K, Trial 2 | -0.002 | -0.172 | 0.084 |
| CT vs. TR-K, Trial 3 | -0.006 | -0.496 | 0.025 |
| CT vs. TR-K, Trial 3 | $-3.625 \times 10^{-6}$ | -0.009 | 0.009 |

### Supplementary Table S8. A summary of measurements in HDG.

| Measurement | Phase (Opponent Strategy) | CT | Theft Recidivist | |
|---|---|---|---|---|
| | | | TR+K | TR-K |
| H-Response | D-phase (Dove) | 11.11 ± 0.91 | 11.77 ± 2.28 | 14.86 ± 1.01 |
| | H-phase (Hawk) | 7.98 ± 0.60 | 10.23 ± 1.98 | 12.34 ± 1.03 |
| Payoffs | D-phase (Dove) | 8.09 ± 0.67 | 10.15 ± 1.02 | 9.33 ± 0.85 |
| | H-Phase (Hawk) | -7.64 ± 1.17 | -11.92 ± 2.52 | -12.70 ± 1.08 |

HDG – Hawk-Dove game; H-Response – The number of aggressive (Hawk) response that participants made; CT – Control subject with no criminal record; TR+K – Theft recidivist with kleptomania; TR-K – Theft recidivist without kleptomania; Payoffs – The net scores for resources that participants obtained (plus) or lost (minus) against the opponents. mean ± s.e.m.

### Supplementary Table S9. Bayesian GLMM for H-response in HDG.

**Posterior Estimate Summaries for Fixed Effects and Interaction Term (Difference from intercept)**

| Variable | Level 1 | Level 2 | Estimate | Est.Error | 95% CI | | $\hat{R}$ | ESS(bulk) | ESS(tail) |
|---|---|---|---|---|---|---|---|---|---|
| | | | | | Lower | Upper | | | |
| Intercept | | | 0.088 | 0.122 | -0.152 | 0.342 | 1.001 | 1870 | 3053 |
| Group | CT | | -0.213 | 0.140 | -0.504 | 0.072 | 1.002 | 1766 | 1834 |
| | TR+K | | -0.012 | 0.209 | -0.430 | 0.425 | 1.001 | 2228 | 3141 |
| | TR-K | | 0.229 | 0.149 | -0.062 | 0.527 | 1.001 | 1879 | 2989 |
| Age | | | 0.133 | 0.091 | -0.053 | 0.313 | 1.002 | 1244 | 2410 |
| Sex | Female | | -0.079 | 0.097 | -0.268 | 0.108 | 1.001 | 1818 | 2510 |
| | Male | | 0.082 | 0.097 | -0.105 | 0.272 | 1.001 | 1807 | 2525 |
| Smoking | No | | -0.124 | 0.114 | -0.341 | 0.099 | 1.003 | 1548 | 2061 |
| | Yes | | 0.127 | 0.115 | -0.097 | 0.344 | 1.003 | 1536 | 1964 |
| Phase | Dove | | -0.018 | 0.038 | -0.091 | 0.056 | 1.001 | 4393 | 4033 |
| | Hawk | | 0.021 | 0.038 | -0.054 | 0.094 | 1.001 | 4429 | 3949 |
| Group x Phase | CT | D-phase | -0.174 | 0.147 | -0.469 | 0.122 | 1.002 | 1846 | 1873 |
| | | H-phase | -0.252 | 0.146 | -0.556 | 0.038 | 1.002 | 1915 | 1813 |
| | TR+K | D-phase | -0.082 | 0.234 | -0.553 | 0.386 | 1.001 | 2332 | 2878 |
| | | H-phase | 0.058 | 0.223 | -0.395 | 0.544 | 1.000 | 2519 | 2835 |
| | TR-K | D-phase | 0.201 | 0.155 | -0.105 | 0.513 | 1.001 | 1894 | 3363 |
| | | H-phase | 0.257 | 0.150 | -0.046 | 0.563 | 1.000 | 2161 | 3454 |

**Estimated marginal means**

| Group | Phase | Median | 95% HPD | |
|---|---|---|---|---|
| | | | Lower | Upper |
| CT | D-phase | -0.088 | -0.372 | 0.222 |
| CT | H-phase | -0.165 | -0.470 | 0.130 |
| TR+K | D-phase | $1.369 \times 10^{-4}$ | -0.631 | 0.684 |
| TR+K | H-phase | 0.139 | -0.485 | 0.815 |
| TR-K | D-phase | 0.289 | 0.028 | 0.549 |
| TR-K | H-phase | 0.343 | 0.079 | 0.602 |

**Pairwise Contrasts**

| Contrasts | Estimate | 95% HPD | |
|---|---|---|---|
| | | Lower | Upper |
| CT vs. TR+K, D-phase | 0.087 | -0.553 | 0.825 |
| CT vs. TR-K, D-phase | 0.376 | -0.038 | 0.778 |
| CT vs. TR+K, H-phase | 0.307 | -0.398 | 1.002 |
| CT vs. TR-K, H-phase | 0.508 | 0.118 | 0.915 |

H-Response – The number of aggressive (Hawk) response that participants made; Est.Error – Estimate error (Posterior standard deviation); $\hat{R}$ – Gelman-Rubin convergence diagnostic; CI – Credible interval; HPD – Highest probability distribution; ESS – Effective sample size; Bulk – Sampling efficiency in the central region of the posterior distribution; Tail – Sampling efficiency in the outer edges of the posterior distribution; D-phase – Trials against the opponent with the dove (cooperative) strategy; H-phase – Trials against the opponent with the hawk (aggressive) strategy.

### Supplementary Table S10. Bayesian GLMM for payoffs in HDG.

**Posterior Estimate Summaries for Fixed Effects and Interaction Term (Difference from intercept)**

| Variable | Level 1 | Level 2 | Estimate | Est.Error | 95% CI | | $\hat{R}$ | ESS(bulk) | ESS(tail) |
|---|---|---|---|---|---|---|---|---|---|
| | | | | | Lower | Upper | | | |
| Intercept | | | -0.008 | 0.087 | -0.188 | 0.164 | 1.002 | 1885 | 712.3 |
| Group | CT | | 0.066 | 0.102 | -0.137 | 0.265 | 1.002 | 3072 | 3091 |
| | TR+K | | -0.006 | 0.143 | -0.298 | 0.270 | 1.001 | 2557 | 917.5 |
| | TR-K | | -0.062 | 0.110 | -0.273 | 0.153 | 1.000 | 3395 | 3107 |
| Age | | | -0.081 | 0.073 | -0.219 | 0.057 | 1.000 | 4642 | 3816 |
| Sex | Female | | -0.016 | 0.079 | -0.174 | 0.137 | 1.001 | 3933 | 3738 |
| | Male | | 0.014 | 0.079 | -0.140 | 0.172 | 1.001 | 3907 | 3679 |
| Smoking | No | | 0.014 | 0.083 | -0.151 | 0.184 | 1.001 | 3680 | 3237 |
| | Yes | | -0.015 | 0.083 | -0.186 | 0.149 | 1.001 | 3694 | 3284 |
| Phase | Dove | | 0.062 | 0.078 | -0.087 | 0.214 | 1.000 | 3577 | 3335 |
| | Hawk | | -0.063 | 0.078 | -0.216 | 0.085 | 1.000 | 3561 | 3407 |
| Group x Phase | CT | D-phase | -0.153 | 0.139 | -0.422 | 0.119 | 1.002 | 3739 | 2170 |
| | | H- phase | 0.284 | 0.138 | 0.019 | 0.554 | 1.000 | 4754 | 4014 |
| | TR+K | D- phase | 0.194 | 0.233 | -0.261 | 0.657 | 1.000 | 2825 | 3239 |
| | | H- phase | -0.207 | 0.242 | -0.690 | 0.249 | 1.003 | 2470 | 2193 |
| | TR-K | D- phase | 0.144 | 0.147 | -0.153 | 0.442 | 1.000 | 2825 | 3239 |
| | | H- phase | -0.267 | 0.149 | -0.564 | 0.039 | 1.000 | 5266 | 3506 |

**Estimated marginal means**

| Group | Phase | Median | 95% HPD | |
|---|---|---|---|---|
| | | | Lower | Upper |
| CT | D-phase | -0.161 | -0.431 | 0.130 |
| CT | H- phase | 0.274 | -0.008 | 0.541 |
| TR+K | D-phase | 0.184 | -0.368 | 0.722 |
| TR+K | H- phase | -0.206 | -0.754 | 0.345 |
| TR-K | D-phase | 0.133 | -0.160 | 0.429 |
| TR-K | H- phase | -0.277 | -0.564 | 0.015 |

**Pairwise Contrasts**

| Contrasts | Estimate | 95% HPD | |
|---|---|---|---|
| | | Lower | Upper |
| CT vs. TR+K, D-phase | 0.350 | -0.264 | 0.914 |
| CT vs. TR-K, D-phase | 0.295 | -0.120 | 0.708 |
| CT vs. TR+K, H- phase | -0.492 | -1.099 | 0.109 |
| CT vs. TR-K, H- phase | -0.550 | -0.966 | -0.158 |
